# Brain Anatomical Covariation Patterns Linked to Binge Drinking and Age at First Full Drink Prior to 21 Years

**DOI:** 10.1101/2020.08.02.232942

**Authors:** Yihong Zhao, R. Todd Constable, Denise Hien, Tammy Chung, Marc N. Potenza

## Abstract

Binge drinking and age at first full drink of alcohol prior to 21 years (AFD<21) have been linked to neuroanatomical differences in cortical and subcortical grey matter (GM) volume, cortical thickness, and surface area. Despite the potential to reveal novel network-level relationships, structural covariation patterns among these morphological measures have yet to be examined relative to binge drinking and AFD<21. Here, we used the Joint and Individual Variance Explained (JIVE) method to characterize structural covariation patterns common across and specific to morphological measures in 293 participants (149 individuals with binge drinking and 144 healthy controls) from the Human Connectome Project (HCP). An independent dataset (Nathan Kline Institute Rockland Sample; NKI-RS) was used to examine reproducibility/ generalizability. We identified a highly reproducible joint component dominated by structural covariation between GM volume in the brainstem and thalamus proper, and GM volume and surface area in prefrontal cortical regions. Using linear mixed regression models, we found that this joint component was related to AFD<21 in both the HCP and NKI-RS datasets, whereas the individual thickness component associated with binge drinking and AFD<21 in the HCP dataset was not statistically significant in the NKI-RS sample. Taken together, our results show that a highly reproducible structural pattern involving covariation in brain regions relevant to thalamic-PFC-brainstem neural circuitry is linked to age at first full drink.

## INTRODUCTION

Age at first full drink prior to 21 years (AFD<21) and subsequent binge drinking (BD) are two important factors linked to the development of alcohol use disorders (AUDs) (1, 2). Adolescence, characterized by imbalanced brain development with subcortical regions developing earlier than the prefrontal control regions, is a critical risk period for addictions (3-5). Early onset of alcohol use may interfere with ongoing neurodevelopment, inducing neurobiological changes that could promote subsequent development of AUDs (1, 2). Human neuroimaging studies indicate that early onset of alcohol use and BD in adolescents or emerging adults are linked to multiple brain structural changes (1, 2), such as reduced cortical thickness (6), decreased surface area (7), and increased grey matter (GM) densities (8) in frontal regions, and a GM volume reduction in many regions (9-11) with exceptions including striatal volumetric increases (12). Sex-related effects have also been observed, with cortical thickness in selected frontal regions thinner in males and thicker in females (13) and putamenal volumes smaller in males and larger in females (14).

Among brain regions related to early onset alcohol use and BD, the prefrontal cortex (PFC), amygdala, and striatum play critical roles in addiction neurocircuitry (1, 15). Early initiation of substance use has been hypothesized to reflect delayed or aberrant development of PFC regions involved in executive functions (1, 4, 5, 15). Recently, pre-clinical research extended models of addiction neurocircuitry (15) to include a focus on the medial PFC-brainstem circuit, demonstrating that neural response in this circuit during initial alcohol exposure predicted the future development of compulsive drinking in a mouse model (16). However, whether such an understudied, yet potentially important, component of addiction neurocircuitry also operates in humans remains to be determined.

Alcohol-use-related structural brain findings to date have been identified mainly via group mean differences (either increased or reduced) in morphological measures in individual brain regions. Such analyses do not typically consider that brain regions function in interacting circuits or networks. Indeed, inter-individual differences in the structure of brain regions within the same or connected neural circuitry often covary more than individual differences in other brain regions, suggesting coordinated development within communities of brain regions (17). Thus, structural covariance analyses may reveal functionally or developmentally linked subsystems. Specifically, structural covariance analyses compare differences in group-level structural covariation networks, where networks are based on pairwise correlations in morphological measures (e.g., cortical thickness) between brain regions. This group-level network-based approach has been used to identify regions suggestive of coordinated development during brain maturation in schizophrenia (18). However, to our knowledge, no published structural covariance studies have examined potentially coordinated development of brain regions in relation to BD or AFD<21.

Here, we sought to investigate whether common and distinct covariation patterns across well-studied cortical morphological measures (thickness, surface area, and GM volume) and subcortical GM volume are associated with BD and/or AFD<21. To accomplish this goal, we used the Joint and Individual Variation Explained (JIVE) method (19). JIVE differs from other network-based structural covariance analyses described above, which are limited to group-level analyses, and thus cannot obtain individual-level measures of structural synchronization. In contrast, JIVE summarizes structural covariation patterns across multiple morphological measures into different component scores. Since brain structures with larger loading magnitudes in a JIVE component are generally more correlated than those with smaller loading magnitudes in the same component (20), the JIVE component scores may provide insight into the extent of synchronized development across brain regions and morphological measures at individual levels. Indeed, our prior work has shown that JIVE can be used to integrate multiple morphological measures into joint and specific components that can robustly predict brain age (20). In short, JIVE analysis may help reveal information at the brain network level, not only providing an efficient data reduction, but also indicating potentially interacting neural circuits.

Given that cortical and subcortical regions are associated with early initiation of alcohol use and BD (1, 2), we hypothesized that cortical and subcortical covariation patterns across multiple morphological measures would be related to AFD<21 and BD. We used the Human Connectome Project (HCP) (21) dataset for our primary analyses and the Nathan Kline Institute-Rockland Sample (NKI-RS (22)) for replication. Our finding of an AFD<21-associated joint component dominated by structural covariation among brainstem, thalamus, and PFC suggests a potential important role of brainstem in alcohol addiction in humans, possibly via the thalamus-PFC-brainstem circuit.

## METHODS AND MATERIALS

### Study Sample

We utilized 293 subjects from the HCP S1200 data (21). The HCP is a community-based sample with participants free of current serious psychiatric conditions or neurologic illness, and free of lifetime substance use disorders (SUDs) other than AUD, cannabis use disorder, and tobacco use disorder. The full HCP S1200 data includes 1,206 participants aged 22-36 years with structural brain imaging data along with alcohol consumption information from the Semi-Structured Assessment for the Genetics of Alcoholism (SSAGA). Our final sample was selected based on the following criteria.

### Inclusion and Exclusion Criteria

For inclusion in analyses, participants met the following criteria: 1) passed image quality controls; 2) had no conflict information between SSAGA survey and laboratory drug results; and, 3) had complete data on SSAGA substance use measures, T1-weighted cortical morphometric features, and potentially confounding variables. Subjects with lifetime AUD status were included only if they reported BD in the past 12 months. Given brain alterations related to use of other substances, subjects with any lifetime SUDs without AUD co-morbidity were excluded from analyses, regardless of past 12-month BD status. Healthy controls (HCs) reported no more than one (for female) or two (for male) drinks in any drinking day in their lifetime. Also exclusionary to HC status were: 1) more than five lifetime uses of any illicit drugs including hallucinogens, opiates, sedatives, and stimulants; and, 2) positive drug tests for methamphetamine, amphetamines, cocaine, opiates, tetrahydrocannabinol, and oxycontin. The final sample consisted of 293 participants (144 HC vs 149 BD).

### BD and Age at First Full Drink Prior to 21 Years (AFD<21)

BD subjects were defined as those who had at least four (for female) or five (for male) drinks within a period of 24 hours at least once per week in the past 12 months. AFD<21 indicated whether the participant had his/her very first full alcoholic drink (e.g., beer, wine, wine coolers, and hard liquor) prior to 21 years. The age of 21 years was chosen as it is the minimum legal drinking age in the US. Per the National Institute on Alcohol Abuse and Alcoholism guidelines, people younger than 21 years should avoid alcohol use completely (https://www.niaaa.nih.gov/alcohol-health/overview-alcohol-consumption/moderate-binge-drinking).

### Validation Data Set

The NKI-RS includes publicly available data (22). We applied similar inclusion and exclusion criteria to select BD and HC subjects. Subjects aged between 18 and 60 years were selected if they met BD criteria in the past year (with or without lifetime AUD diagnosis). HC subjects were required to 1) be free of any SUDs, and 2) have no self-reported BD history, limited alcohol use, and no or only occasional past-year use of other substances. The selected sample consisted of 93 subjects (46 HC and 47 BD). Potentially confounding variables were controlled in regression analyses, including age, sex, race (white vs. other), socioeconomic status, fluid intelligence (Wechsler Abbreviated Scale of Intelligence-II), handedness, and estimated total intracranial volume.

### Brain Image Processing

All T1-weighted imaging data in the HCP sample were acquired on a customized Siemens 3T Skyra scanner using a multi-band sequence at a spatial resolution of 0.7 mm isotropic voxels (21). Structural scans were preprocessed using the customized HCP structural pipeline based on FreeSurfer 5.3 and quality controlled by the HCP team. The tabulated structural dataset includes multiple morphometric measures for each subject. Based on prior findings from studies of early initiation of alcohol use and BD, we investigated structural covariation patterns among GM volumes in 17 subcortical regions (left and right amygdala, accumbens area, caudate, hippocampus, putamen, pallidum, thalamus proper, ventral diencephalon, and brainstem), and cortical thickness, surface area and GM volume in 68 regions-of-interest (ROIs) based on the Desikan-Killiany atlas (23).

Subjects in the NKI-RS sample underwent a scan session using a Siemens TrioTM 3.0T MRI scanner. T1-weighted images were acquired using a magnetization-prepared rapid gradient echo (MPRAGE) sequence with 1mm isotropic resolution. The structural images were preprocessed using the recon-all pipeline from FreeSurfer version 5.3.0 (24, 25), an extensively used robust pipeline optimized for 1mm isotropic data. Image pre-processing quality was checked following ENIGMA image quality control protocols (http://enigma.ini.usc.edu/protocols/imaging-protocols/).

### Statistical Analysis of Cortical and Subcortical Covariation Patterns

Subcortical GM volume, cortical thickness, surface area, and cortical GM volume were treated as four data sources. We utilized JIVE (19) to identify covariation patterns consistent across different morphological measures and patterns unique to individual morphological measures. JIVE is a dimension-reduction and pattern-discovery approach for data from multiple resources. Specifically, JIVE decomposes total variation into three terms: joint variation across multiple morphometric measures, structured variation unique to each morphometric measure, and residual noise to be discarded from analyses.

Briefly, let *X*_1,_ *X*_2,_*X*_3_, *X*_4_ be data matrices for four morphological measures in which each row stands for a morphometric feature (e.g., surface area, cortical thickness) and each column for a subject, *A*_*i*_ be matrix for individual structure of *X*_*i*_, *J*_*i*_ be the submatrix of joint structure matrix associated with *X*_*i*_, and *∈*_*i*_ be error matrix of *X*_*i*_. The JIVE model can be written as *X*_1_ = J_1_ + *A*_1_ + ∈_1_, …, *X* _4_ = *J*_4_ + *A*_4_ + *∈*_*4*_. An iterative approach was used to estimate the loadings of *J*_*i*_ and *A*_*i*_ (i.e., joint and individual components). The optimal number of joint and individual components was determined via a permutation approach (19) with 10,000 permutations and the significance level set to 0.0001. All procedures were implemented in R using modified functions from the r.jive package.

### Linear Mixed Regression Analysis

To control for potential confounding effects in regression analysis, we included covariates of age, sex, race (white vs. other), education and income level (to approximate socioeconomic status), twin status (monozygotic, dizygotic, or unrelated), fluid intelligence score based on Raven’s Progressive Matrices, and estimated total intracranial volume (to control for differences in overall head size). Linear mixed models with family as a random effect were used to test whether JIVE joint and individual components were related to BD status and AFD<21. To control for multiple comparisons, the false discovery rate (FDR) was controlled at 0.05 significance.

## RESULTS

### Participant Characteristics (HCP Sample)

The HCP sample included 144 HCs, 61 BD subjects without lifetime AUD diagnosis (BD-AUD), and 88 BD subjects with lifetime AUD diagnosis (BD+AUD) (Table 1). These groups did not differ significantly on age, handedness, household income, education, intelligence, zygosity, and family history of drug or alcohol use. However, our bivariate analyses indicated that there were significantly more white males in the BD group. Also, the HC group on average had significantly smaller estimated intracranial volumes than the BD group. As expected, there was a significant difference in age at first use of alcohol. The mean age at first full drink was 20.3 (±2.8) years for HC, 17.1 (±2.5) years for BD-AUD, and 16.0 (±1.9) years for BD+AUD.

**Table 1.**
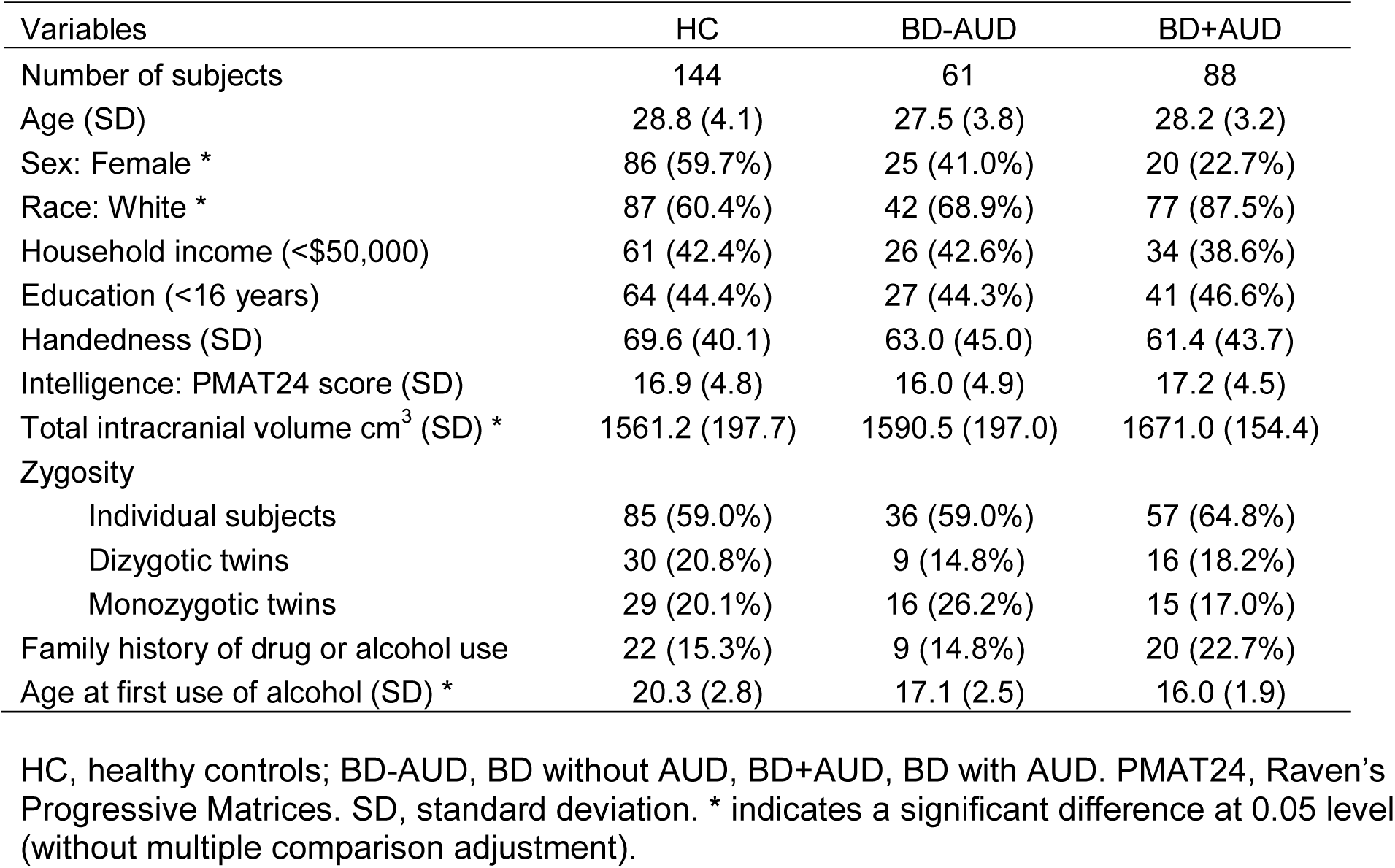
Summary of binge drinking and heathy control subjects from the HCP sample used in this study.

| Variables | HC | BD-AUD | BD+AUD |
| --- | --- | --- | --- |
| Number of subjects | 144 | 61 | 88 |
| Age (SD) | 28.8 (4.1) | 27.5 (3.8) | 28.2 (3.2) |
| Sex: Female * | 86 (59.7%) | 25 (41.0%) | 20 (22.7%) |
| Race: White * | 87 (60.4%) | 42 (68.9%) | 77 (87.5%) |
| Household income (<\$50,000) | 61 (42.4%) | 26 (42.6%) | 34 (38.6%) |
| Education (<16 years) | 64 (44.4%) | 27 (44.3%) | 41 (46.6%) |
| Handedness (SD) | 69.6 (40.1) | 63.0 (45.0) | 61.4 (43.7) |
| Intelligence: PMAT24 score (SD) | 16.9 (4.8) | 16.0 (4.9) | 17.2 (4.5) |
| Total intracranial volume cm<sup>3</sup> (SD) * | 1561.2 (197.7) | 1590.5 (197.0) | 1671.0 (154.4) |
| Zygosity |  |  |  |
| Individual subjects | 85 (59.0%) | 36 (59.0%) | 57 (64.8%) |
| Dizygotic twins | 30 (20.8%) | 9 (14.8%) | 16 (18.2%) |
| Monozygotic twins | 29 (20.1%) | 16 (26.2%) | 15 (17.0%) |
| Family history of drug or alcohol use | 22 (15.3%) | 9 (14.8%) | 20 (22.7%) |
| Age at first use of alcohol (SD) * | 20.3 (2.8) | 17.1 (2.5) | 16.0 (1.9) |

HC, healthy controls; BD-AUD, BD without AUD, BD+AUD, BD with AUD. PMAT24, Raven's Progressive Matrices. SD, standard deviation. \* indicates a significant difference at 0.05 level (without multiple comparison adjustment).

### Cortical and Subcortical Covariation Patterns

JIVE analysis of 221 brain features (three cortical morphological measures for each of 68 ROIs plus 17 subcortical GM volumes) from the HCP sample led to identification of 14 brain signatures (i.e., components). These included one joint component, and three, four, four, and two individual components specific to surface area, cortical thickness, and GM volumes in cortical regions and GM volume in subcortical regions, respectively. Figure 1 shows joint and individual variation across the four morphological measures. Overall, the joint component explained 38.4% of total variation, and the individual components collectively explained 27.6% of total variation. A considerable amount of residual noise was identified in cortical thickness (48.6%), surface area (37.3%), and GM volume (36.0%).

**Fig. 1.**
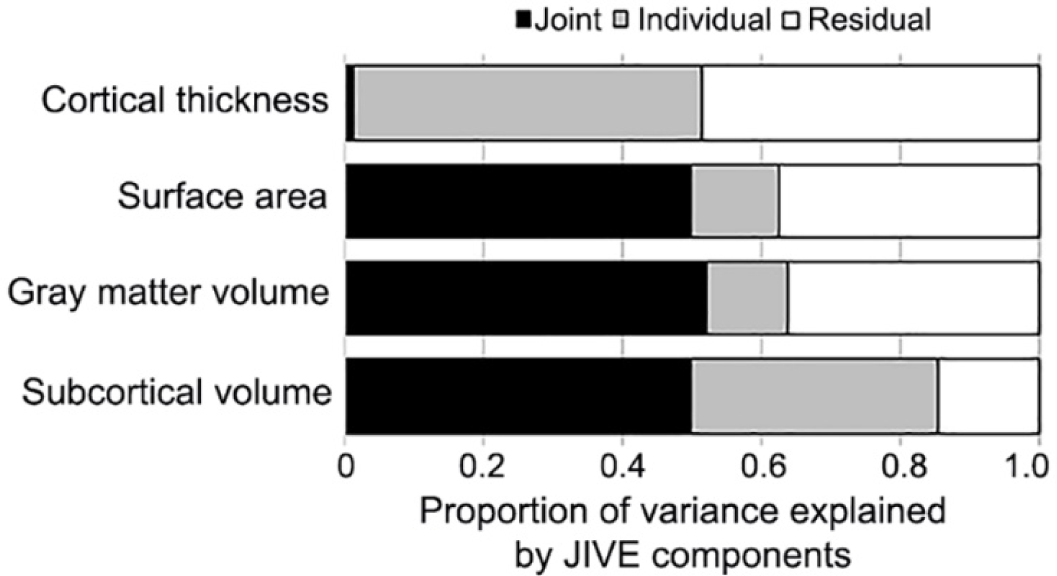
Proportion of variance explained by JIVE components in the HCP data.

### Wide-Spread Cortical Thinning Associated with BD

Controlling for confounding variables, separate linear mixed models were used to test associations between each JIVE component and BD status. After multiple-comparison adjustment, one individual component specific to cortical thickness was negatively associated with BD status, suggesting that past-12-month BD was linked to smaller mean cortical thickness component scores in both BD-AUD (beta=-0.062, p-value=0.001) and BD+AUD (beta=-0.057, p-value=0.003) groups (Figure 2A). Post-hoc subgroup analyses revealed no differences in mean thickness component scores between BD-AUD and BD+AUD groups (beta=0.013, p-value=0.544). To facilitate the interpretation of this cortical thickness component, we listed loadings for each of the 68 ROIs (Table S1). Loadings in most ROIs (∼72%) ranged between 0.1 and 0.15. Seven ROIs (mainly in the temporal lobe: right entorhinal, left and right temporal pole, transverse temporal region, and frontal pole) had loadings between 0.15 and 0.21, suggesting that BD might have a slightly stronger relationship with systematic cortical thinning in temporal lobe regions compared to others. Overall, results suggest that wide-spread cortical thinning is associated with past-12-month BD.

**Figure 2.**
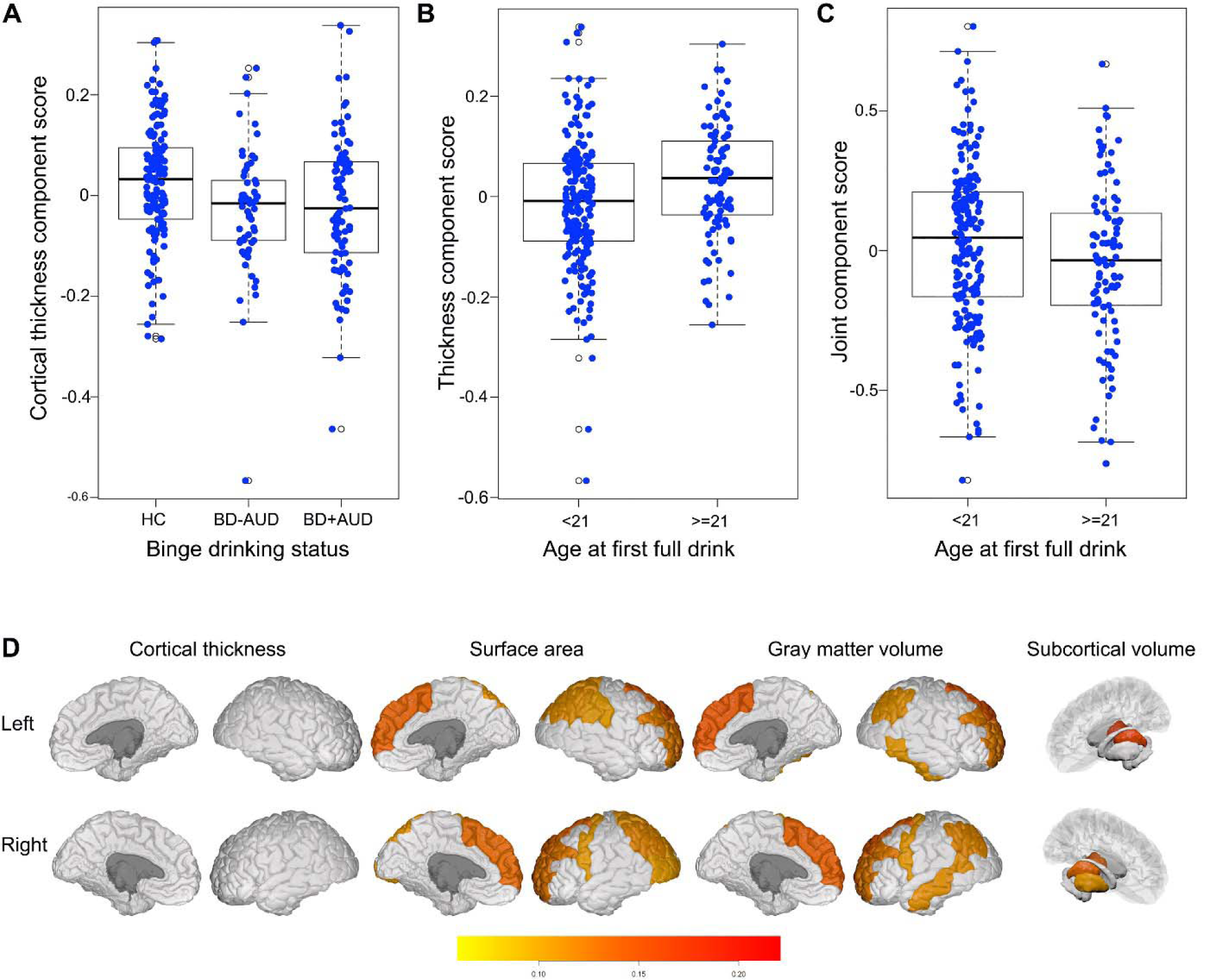
JIVE components related to binge-drinking and age at first full drink prior to 21 years in the HCP data. (**A** & **B**) Boxplots showing a cortical thickness component related to binge-drinking status (p-value=0.003) and to age at first alcohol use (p< 0.001). (**C**) Boxplot showing the JIVE joint component related to age at first full drink (p=0.001). (**D**) Plots of brain regions in the joint component with loadings larger than 0.10. Interior and exterior views of the brain regions are presented for each morphological measure. Brainstem is not shown. HC, healthy controls; BD-AUD, subjects with BD in the past 12 months but without AUD diagnosis; BD+AUD, subjects with both BD in the past 12 months and AUD diagnosis.

### Brain Anatomical Covariation Patterns Associated with AFD<21

Separate linear mixed models were used to test associations between JIVE components and AFD<21, controlling for potentially confounding variables. After multiple-comparison adjustment, two components were associated with AFD<21. First, the aforementioned cortical thickness component reflecting wide-spread cortical thinning associated with BD (Figure 2A) was also related to AFD<21 (beta=-0.066, p-value<0.001; Figure 2B). Second, individuals with AFD<21 displayed greater mean joint component scores (beta=0.059, p-value=0.001; Figure 2C). This joint component, which includes structural covariation pattern across all 221 cortical and subcortical brain features considered (Table S2), is dominated by covariation patterns among subcortical and cortical GM volumes and cortical surface areas, and accounts for roughly 50% of variation within each of these three morphological measures. This joint component, however, accounted for only 1.3% of variation in cortical thickness. Most brain features (147 out of 221, 66.5%) had loading magnitudes less than 0.05, 16 (7.2%) features had loadings between 0.05 and 0.08, 34 (15.4%) had loadings between 0.08 and 0.10, and 24 brain features (10.9%) had loadings greater than 0.10. The 24 brain features with the largest loadings and their individual associations with AFD<21 are listed (Table 2), and the loading plot of these regions is shown (Figure 2D). Figure S1 shows regions with loadings larger than 0.08. Among all 221 brain features, the brainstem had the largest loading (0.433), followed by the thalamus proper (left and right) and PFC regions (e.g., GM volume and surface area in left and right superior frontal and rostral middle frontal regions), suggesting that the joint component is dominated by synchronized/coordinated development in these brain regions. Surprisingly, most of these regions were not significantly related to AFD<21 based on the mixed model regression analysis (Table 2, and Table S2). Thus, our analysis suggests that AFD<21 may not lead to substantial individual brain alterations. Instead, AFD<21 may modify coordinated development among morphological measures in regions including brainstem, thalamus proper, and PFC, although longitudinal studies are needed to investigate this possibility. In addition, several cortical regions in the parietal, temporal and occipital lobes, and other subcortical regions including left and right putamen, hippocampus, caudate, and ventral diencephalon also had relatively large loadings in the joint component (Table S2).

**Table 2.**
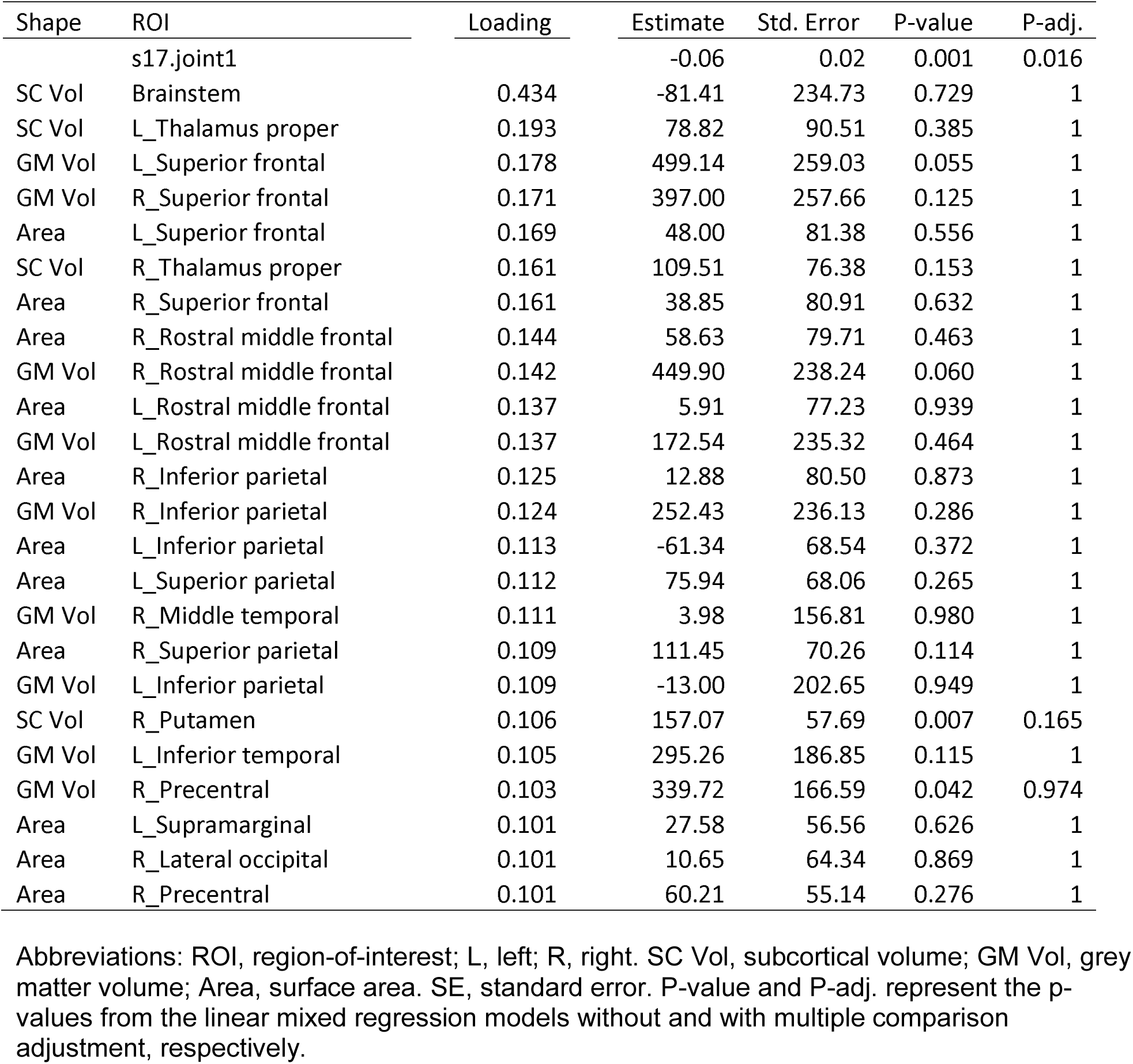
Loading of top brain regions in the joint component related to AFD<21. A total of 24 brain regions and morphological measures in the joint component showing a loading of larger than 0.100 are listed based on the order of the magnitudes of the loadings. The results from individual linear mixed regression analyses are also listed.

| Shape | ROI | Loading | Estimate | Std. Error | P-value | P-adj. |
| --- | --- | --- | --- | --- | --- | --- |
|  | s17.joint1 |  | -0.06 | 0.02 | 0.001 | 0.016 |
| SC Vol | Brainstem | 0.434 | -81.41 | 234.73 | 0.729 | 1 |
| SC Vol | L_Thalamus proper | 0.193 | 78.82 | 90.51 | 0.385 | 1 |
| GM Vol | L_Superior frontal | 0.178 | 499.14 | 259.03 | 0.055 | 1 |
| GM Vol | R_Superior frontal | 0.171 | 397.00 | 257.66 | 0.125 | 1 |
| Area | L_Superior frontal | 0.169 | 48.00 | 81.38 | 0.556 | 1 |
| SC Vol | R_Thalamus proper | 0.161 | 109.51 | 76.38 | 0.153 | 1 |
| Area | R_Superior frontal | 0.161 | 38.85 | 80.91 | 0.632 | 1 |
| Area | R_Rostral middle frontal | 0.144 | 58.63 | 79.71 | 0.463 | 1 |
| GM Vol | R_Rostral middle frontal | 0.142 | 449.90 | 238.24 | 0.060 | 1 |
| Area | L_Rostral middle frontal | 0.137 | 5.91 | 77.23 | 0.939 | 1 |
| GM Vol | L_Rostral middle frontal | 0.137 | 172.54 | 235.32 | 0.464 | 1 |
| Area | R_Inferior parietal | 0.125 | 12.88 | 80.50 | 0.873 | 1 |
| GM Vol | R_Inferior parietal | 0.124 | 252.43 | 236.13 | 0.286 | 1 |
| Area | L_Inferior parietal | 0.113 | -61.34 | 68.54 | 0.372 | 1 |
| Area | L_Superior parietal | 0.112 | 75.94 | 68.06 | 0.265 | 1 |
| GM Vol | R_Middle temporal | 0.111 | 3.98 | 156.81 | 0.980 | 1 |
| Area | R_Superior parietal | 0.109 | 111.45 | 70.26 | 0.114 | 1 |
| GM Vol | L_Inferior parietal | 0.109 | -13.00 | 202.65 | 0.949 | 1 |
| SC Vol | R_Putamen | 0.106 | 157.07 | 57.69 | 0.007 | 0.165 |
| GM Vol | L_Inferior temporal | 0.105 | 295.26 | 186.85 | 0.115 | 1 |
| GM Vol | R_Precentral | 0.103 | 339.72 | 166.59 | 0.042 | 0.974 |
| Area | L_Supramarginal | 0.101 | 27.58 | 56.56 | 0.626 | 1 |
| Area | R_Lateral occipital | 0.101 | 10.65 | 64.34 | 0.869 | 1 |
| Area | R_Precentral | 0.101 | 60.21 | 55.14 | 0.276 | 1 |

Abbreviations: ROI, region-of-interest; L, left; R, right. SC Vol, subcortical volume; GM Vol, grey matter volume; Area, surface area. SE, standard error. P-value and P-adj. represent the p-values from the linear mixed regression models without and with multiple comparison adjustment, respectively.

We also assessed whether age and sex were related to the joint component. Without controlling for confounding variables, the joint component was inversely related to biological age (beta=-0.011, p-value=0.001), and males had larger joint component scores (beta=0.311, p-value<0.001). Controlling for race, socioeconomic status, handedness, and estimated intracranial volumes, relationships with age (beta=-0.006, p-value=0.013) and sex (beta=0.064, p-value=0.003) remained significant.

### Examining NKI-RS Data

To investigate replication in an independent dataset, we conducted JIVE analyses in the NKI-RS sample (Table S3). First, proportions of joint and individual variance explained were highly similar in the NKI-RS (Figure 3A) and HCP (Figure 1) samples. Second, the joint component identified in HCP analyses was highly reproducible in NKI-RS analyses. Pairwise Pearson correlation coefficients between the component loadings from both datasets were used to assess degrees of similarity in brain components. If an HCP component was correlated highly with more than one NKI-RS component, the maximal correlation coefficient was reported. We found that the HCP and NKI-RS joint components correlated highly (r=0.959; Figure 3B). Third, three individual components were also reproducible including components specific to subcortical GM volume (r=0.990), surface area (r=0.881), and cortical thickness (r=0.819), respectively. Correlations in loadings for other components specific to individual morphological measures were low to moderate (ranging from 0.234 to 0.743), indicating that these components were less reproducible. Of note, the thickness component related to BD (Figure 2A) and AFD<21 (Figure 2B) identified from the HCP sample had moderate reproducibility in the NKI-RS sample, as evidenced by the maximal correlation of 0.624 between loadings of the cortical thickness component in HCP and NKI-RS samples.

**Fig. 3.**
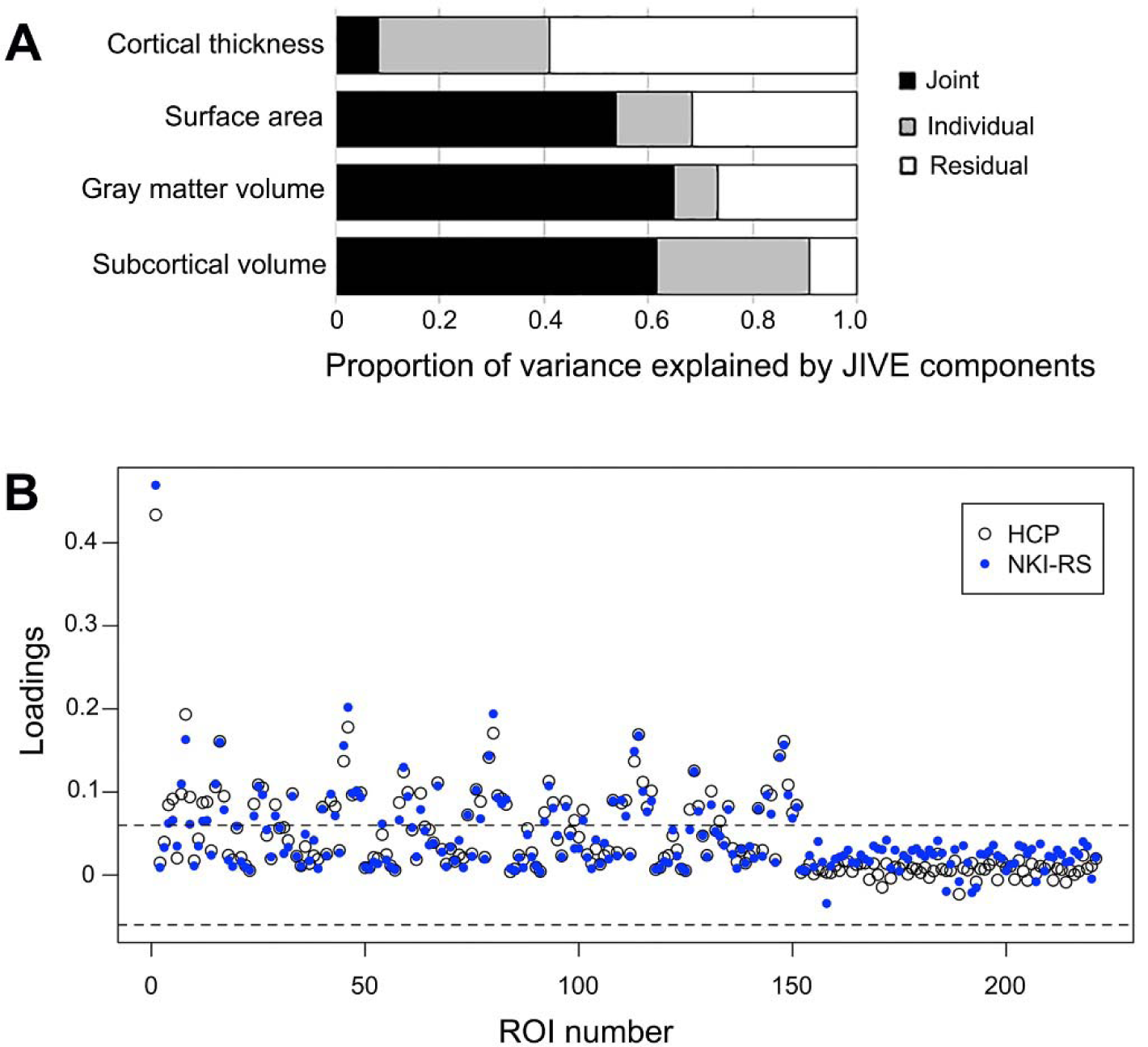
A highly similar joint component exists in the NKI-RS data (shown) as identified in the HCP data. (**A**) Proportion of variance explained by JIVE components in the NKI-RS data. See Figure 1 for proportion of variance explained by JIVE components in the HCP data. (**B**) Correlation of the loadings for 221 ROI structural features in the joint components derived from the HCP and the NKI-RS data sets.

We also investigated whether relationships between two components (i.e., joint and thickness) and BD and/or AFD<21 were reproducible. Regression analysis, controlling for potential confounding effects, showed that neither BD (p-value=0.406) nor AFD<21 (p-value=0.743) were significantly related to the thickness component in the NKI-RS. In contrast, the NKI-RS sample replicated the positive association between the joint component and AFD<21 (beta=0.023, p-value=0.040).

In terms of age and sex, the joint component was inversely related to biological age in NKI-RS with (beta=-0.040, p-value<0.001) or without (beta=-0.032, p-value<0.001) controlling for confounding variables. The significant relationship between sex and the joint component was observed without controlling for confounding variables (beta=1.019, p-value<0.001), but no longer remained (beta=0.014, p-value=0.135) if controlling for confounding variables.

## DISCUSSION

### Joint Structural Covariation Implicates Thalamus-PFC-Brainstem Circuitry

Our hypotheses were largely supported in that we observed joint structural covariation suggestive of widespread cortical thinning related to BD and AFD<21 in HCP analyses and highly correlated joint structural covariation patterns linked to AFD<21 across HCP and NKI-RS datasets. Thus, the latter, replicable results may be more generalizable across samples. Although the former results resonate with those assessing relations between cortical thickness and alcohol misuse using HCP data and different analytical approaches (26), we did not observe these associations in the NKI-RS sample. Therefore, interpretation of relations between the JIVE thickness component and BD and AFD<21 should be made cautiously.

A particularly consistent finding across samples is the association of a cortical and subcortical covariation pattern, or joint component, with AFD<21. The joint component is dominated by covariation between GM volume and surface area features across multiple regions (Table 2). Among all brain features, covariation among brainstem, thalamus proper, and superior and rostral middle frontal cortices contributed most. The PFC has been shown to play a critical role in early alcohol use initiation, as it is still maturing in adolescence and thus may not have sufficient control over sensation- or novelty-seeking which is promoted by subcortical regions and peaks in adolescence, according to dual-systems or maturational imbalance models (1, 4, 5). Although much less is known regarding the role of brainstem in youth alcohol use, alcohol use has been linked to impairment of lower-level brainstem functioning (27), and adolescents who initiated heavy drinking showed reduced brainstem volume (9). Importantly, the JIVE brainstem result appears to be consistent with a recent pre-clinical optogenetic study, which found that neural response in medial-PFC-brainstem circuitry during initial alcohol exposure in a mouse model predicted future compulsive drinking (16). However, the extent to which the PFC-brainstem circuitry contributes to human alcohol use remains unknown. Our JIVE finding that the brainstem contributes most strongly to the AFD<21-related joint component and covaries with PFC regions suggests a key role of brainstem in human alcohol addiction, likely via PFC-brainstem circuitry.

Additionally, the identified frontal and brainstem regions also covary with thalamic GM volumes. The thalamus has been described as a “passive” information relay station, but data suggest that it contributes importantly to cognition (28). Thalamic function via corticothalamic or thalamocortical pathways integrates inputs from the PFC and other cortices. Indeed, the PFC and thalamus can be activated by alcohol cues (29), and decreased connectivity in the thalamus-PFC fibre pathway is associated with AUD (30). In addition, a thalamus-dorsomedial-PFC circuit was recently demonstrated to link to social dominance in mice (31). Notably, social behaviors also have been associated with PFC-brainstem circuitry in the mouse model (32), further suggesting coherence in the coordinated activities of these structures in the thalamus-PFC-brainstem circuit. Given these data and our finding that the brainstem, thalamus and frontal regions are top contributors to the joint component score, we postulate a role for a thalamus-PFC-brainstem circuit in early alcohol use initiation in humans.

Other cortical and subcortical structures also showing strong contributions to the joint component have been implicated in early alcohol use initiation. For example, differences in GM volume in parietal, temporal and occipital lobes have been reported in prior studies examining brain structural changes related to pre-initiation or post-drinking (9, 10, 33, 34). These regions are associated with various functions, such as motor control, emotion processing, language comprehension, and visuospatial processing. Several subcortical regions are noteworthy, including putamen, caudate and their functionally connected regions including the hippocampus and structures in the ventral diencephalon. The dorsal striatum, including putamen and caudate, has been implicated in addiction processes, particularly with respect to habitual versus goal-directed behaviors (35-37). Consistently, the dorsal striatum is proposed as contributing importantly to binge/intoxication phases of AUDs (15). Hippocampal volumetric differences have been reported in adolescents with and without AUDs (38), hippocampus-dorsal-striatum connections are involved in formation of memories and habits may contribute (39), and the hippocampus-thalamus fibre pathway is related to AUD (30). The ventral diencephalon includes structures implicated in controlling alcohol consumption, such as the substantia nigra, subthalamic nucleus and hypothalamus (40-44). In summary, given extensive interconnections between the top three regions (thalamus, PFC, and brainstem) and other cortical or subcortical structures, these latter regions and related circuits may provide or modulate inputs to or outputs from the major thalamus-PFC-brainstem circuit in balancing impulse control vs. sensation- or novelty-seeking that may influence AFD<21. Cortical and subcortical covariation patterns may represent a potential brain-based biomarker for AFD<21.

### Structural Covariation May Reflect Synchronized Development

A major advantage of JIVE analysis is that it can effectively consolidate covariation patterns among different morphological measures into lower dimensional representations (i.e., component scores). This is critically important for generating meaningful, interpretable results using existing statistical models (45, 46). Other methods, such as structural learning and integrative decomposition of multi-view data (47), common orthogonal basis extraction (48), and group factor analysis (49), may also be used. Our results suggest that an attractive feature of JIVE is the performance robustness, consistent with our prior study of brain age prediction (20). Given that brain morphological measures within structurally and functionally connected regions often co-vary (17), the identified JIVE covariation patterns may be interpreted as synchronized or coordinated development of brain structures across cortical and subcortical regions, although longitudinal studies should examine this directly. Longitudinal studies may also disentangle the extent to which such relationships exist prior to or subsequent to alcohol use. Nonetheless, our study underscores the importance of using innovative data-driven approaches to extract novel information from big data to provide new insight into alcohol-use behaviors/problems.

### Limitations

Our work is cross-sectional and retrospective; thus, it is not possible to determine whether this joint structural covariation pattern reflects a consequence or a precursor of AFD<21. Also, as most HCP participants initiated alcohol use prior to tobacco and/or cannabis use, it remains unclear whether the identified structural covariation pattern is specific to alcohol use. Future studies using longitudinal data are needed to determine: 1) whether the identified structural covariation pattern can prospectively predict initiation of use of alcohol and/or other substances; 2) how developmental trajectories of brain structural covariation patterns change from childhood to adolescence; and, 3) whether this structural covariation pattern predicts risk or resilience relative to substance use. Additionally, due to limited sample sizes, sex, age and other factors were treated as covariates. Future studies using JIVE in larger samples may provide insights into effects of these individual differences on structural covariation patterns and their relationship to BD and AFD<21.

### Conclusions

JIVE identified a highly reproducible cortical and subcortical structural covariation pattern involving brain regions relevant to thalamic-PFC-brainstem neural circuitry. Further, a covariation pattern was linked to AFD<21 in both HCP and NKI-RS datasets. This data-driven discovery study highlights the importance of considering subcortical and cortical regions together to increase understanding of AFD<21 correlates.

## ACKNOWLEDGEMENTS

This work was in part funded by National Institutes of Health R21AA023800 (Zhao), R01DA039136 (Potenza), and R01AA025853 (Hien). Data were provided by the Human Connectome Project, WU-MINN Consortium (Principal Investigators: David Van Essen and Kamil Ugurbil; 1U54MH091657) funded by the 16 NIH institutes and centers that support the NIH Blueprint for Neuroscience Research; and by the McDonnell Center for Systems Neuroscience at Washington University.

## SUPPLEMENTAL INFORMATION

**Table S1.** Loading of brain regions in the individual thickness component related to binge drinking and AFD<21 in the HCP sample.

**Table S2.** Loading of all 221 brain features in the joint component related to AFD<21 in the HCP sample.

**Table S3.** Summary of binge drinking and heathy control subjects from the NKI-RS sample used in this study.

**Figure S1.** Brain regions in the joint component with loadings larger than 0.08.

## REFERENCES

1. Squeglia LM, Cservenka A (2017): Adolescence and Drug Use Vulnerability: Findings from Neuroimaging. Current opinion in behavioral sciences. 13:164–170.

2. Lees B, Meredith LR, Kirkland AE, Bryant BE, Squeglia LM (2020): Effect of alcohol use on the adolescent brain and behavior. Pharmacology, biochemistry, and behavior. 192:172906.

3. Chambers RA, Taylor JR, Potenza MN (2003): Developmental neurocircuitry of motivation in adolescence: a critical period of addiction vulnerability. The American journal of psychiatry. 160:1041–1052.

4. Casey B, Jones RM, Somerville LH (2011): Braking and Accelerating of the Adolescent Brain. J Res Adolesc. 21:21–33.

5. Shulman EP, Smith AR, Silva K, Icenogle G, Duell N, Chein J, et al. (2016): The dual systems model: Review, reappraisal, and reaffirmation. Developmental cognitive neuroscience. 17:103–117.

6. Brumback TY, Worley M, Nguyen-Louie TT, Squeglia LM, Jacobus J, Tapert SF (2016): Neural predictors of alcohol use and psychopathology symptoms in adolescents. Development and psychopathology. 28:1209–1216.

7. Infante MA, Courtney KE, Castro N, Squeglia LM, Jacobus J (2018): Adolescent Brain Surface Area Pre- and Post-Cannabis and Alcohol Initiation. Journal of studies on alcohol and drugs. 79:835–843.

8. Sousa SS, Sampaio A, Marques P, Goncalves OF, Crego A (2017): Gray Matter Abnormalities in the Inhibitory Circuitry of Young Binge Drinkers: A Voxel-Based Morphometry Study. Frontiers in psychology. 8:1567.

9. Squeglia LM, Rinker DA, Bartsch H, Castro N, Chung Y, Dale AM, et al. (2014): Brain volume reductions in adolescent heavy drinkers. Developmental cognitive neuroscience. 9:117–125.

10. Yang X, Tian F, Zhang H, Zeng J, Chen T, Wang S, et al. (2016): Cortical and subcortical gray matter shrinkage in alcohol-use disorders: a voxel-based meta-analysis. Neuroscience and biobehavioral reviews. 66:92–103.

11. Baranger DAA, Demers CH, Elsayed NM, Knodt AR, Radtke SR, Desmarais A, et al. (2020): Convergent Evidence for Predispositional Effects of Brain Gray Matter Volume on Alcohol Consumption. Biological psychiatry. 87:645–655.

12. Howell NA, Worbe Y, Lange I, Tait R, Irvine M, Banca P, et al. (2013): Increased ventral striatal volume in college-aged binge drinkers. PloS one. 8:e74164.

13. Squeglia LM, Sorg SF, Schweinsburg AD, Wetherill RR, Pulido C, Tapert SF (2012): Binge drinking differentially affects adolescent male and female brain morphometry. Psychopharmacology. 220:529–539.

14. Fein G, Greenstein D, Cardenas VA, Cuzen NL, Fouche JP, Ferrett H, et al. (2013): Cortical and subcortical volumes in adolescents with alcohol dependence but without substance or psychiatric comorbidities. Psychiatry research. 214:1–8.

15. Koob GF, Volkow ND (2016): Neurobiology of addiction: a neurocircuitry analysis. Lancet Psychiatry. 3:760–773.

16. Siciliano CA, Noamany H, Chang CJ, Brown AR, Chen X, Leible D, et al. (2019): A cortical-brainstem circuit predicts and governs compulsive alcohol drinking. Science. 366:1008–1012.

17. Alexander-Bloch A, Giedd JN, Bullmore E (2013): Imaging structural co-variance between human brain regions. Nature reviews Neuroscience. 14:322–336.

18. Sandini C, Scariati E, Padula MC, Schneider M, Schaer M, Van De Ville D, et al. (2018): Cortical Dysconnectivity Measured by Structural Covariance Is Associated With the Presence of Psychotic Symptoms in 22q11.2 Deletion Syndrome. Biological psychiatry Cognitive neuroscience and neuroimaging. 3:433–442.

19. Lock EF, Hoadley KA, Marron JS, Nobel AB (2013): Joint and Individual Variation Explained (Jive) for Integrated Analysis of Multiple Data Types. The annals of applied statistics. 7:523–542.

20. Zhao Y, Klein A, Castellanos FX, Milham MP (2019): Brain age prediction: Cortical and subcortical shape covariation in the developing human brain. NeuroImage. 202:116149.

21. Van Essen DC, Smith SM, Barch DM, Behrens TE, Yacoub E, Ugurbil K, et al. (2013): The WU-Minn Human Connectome Project: an overview. NeuroImage. 80:62–79.

22. Nooner KB, Colcombe SJ, Tobe RH, Mennes M, Benedict MM, Moreno AL, et al. (2012): The NKI-Rockland Sample: A Model for Accelerating the Pace of Discovery Science in Psychiatry. Frontiers in neuroscience. 6:152.

23. Desikan RS, Segonne F, Fischl B, Quinn BT, Dickerson BC, Blacker D, et al. (2006): An automated labeling system for subdividing the human cerebral cortex on MRI scans into gyral based regions of interest. NeuroImage. 31:968–980.

24. Fischl B, Salat DH, Busa E, Albert M, Dieterich M, Haselgrove C, et al. (2002): Whole brain segmentation: automated labeling of neuroanatomical structures in the human brain. Neuron. 33:341–355.

25. Fischl B, van der Kouwe A, Destrieux C, Halgren E, Segonne F, Salat DH, et al. (2004): Automatically parcellating the human cerebral cortex. Cerebral cortex. 14:11–22.

26. Morris VL, Owens MM, Syan SK, Petker TD, Sweet LH, Oshri A, et al. (2019): Associations Between Drinking and Cortical Thickness in Younger Adult Drinkers: Findings From the Human Connectome Project. Alcoholism, clinical and experimental research. 43:1918–1927.

27. Oscar-Berman M, Marinkovic K (2003): Alcoholism and the brain: an overview. Alcohol research & health : the journal of the National Institute on Alcohol Abuse and Alcoholism. 27:125–133.

28. Wolff M, Vann SD (2019): The Cognitive Thalamus as a Gateway to Mental Representations. The Journal of neuroscience : the official journal of the Society for Neuroscience. 39:3–14.

29. George MS, Anton RF, Bloomer C, Teneback C, Drobes DJ, Lorberbaum JP, et al. (2001): Activation of prefrontal cortex and anterior thalamus in alcoholic subjects on exposure to alcohol-specific cues. Archives of general psychiatry. 58:345–352.

30. Segobin S, Laniepce A, Ritz L, Lannuzel C, Boudehent C, Cabe N, et al. (2019): Dissociating thalamic alterations in alcohol use disorder defines specificity of Korsakoff’s syndrome. Brain : a journal of neurology. 142:1458–1470.

31. Zhou T, Zhu H, Fan Z, Wang F, Chen Y, Liang H, et al. (2017): History of winning remodels thalamo-PFC circuit to reinforce social dominance. Science. 357:162–168.

32. Franklin TB, Silva BA, Perova Z, Marrone L, Masferrer ME, Zhan Y, et al. (2017): Prefrontal cortical control of a brainstem social behavior circuit. Nature neuroscience. 20:260–270.

33. van Holst RJ, de Ruiter MB, van den Brink W, Veltman DJ, Goudriaan AE (2012): A voxel-based morphometry study comparing problem gamblers, alcohol abusers, and healthy controls. Drug and alcohol dependence. 124:142–148.

34. Segobin SH, Chetelat G, Le Berre AP, Lannuzel C, Boudehent C, Vabret F, et al. (2014): Relationship between brain volumetric changes and interim drinking at six months in alcohol-dependent patients. Alcoholism, clinical and experimental research. 38:739–748.

35. Chen G, Cuzon Carlson VC, Wang J, Beck A, Heinz A, Ron D, et al. (2011): Striatal involvement in human alcoholism and alcohol consumption, and withdrawal in animal models. Alcoholism, clinical and experimental research. 35:1739–1748.

36. Haber SN (2016): Corticostriatal circuitry. Dialogues in clinical neuroscience. 18:7–21.

37. Cox J, Witten IB (2019): Striatal circuits for reward learning and decision-making. Nature reviews Neuroscience. 20:482–494.

38. De Bellis MD, Clark DB, Beers SR, Soloff PH, Boring AM, Hall J, et al. (2000): Hippocampal volume in adolescent-onset alcohol use disorders. The American journal of psychiatry. 157:737–744.

39. Volkow ND, Morales M (2015): The Brain on Drugs: From Reward to Addiction. Cell. 162:712–725.

40. Morales-Mulia M, de Gortari P, Amaya MI, Mendez M (2013): Acute ethanol administration differentially alters enkephalinase and aminopeptidase N activity and mRNA levels in regions of the nigrostriatal pathway. J Mol Neurosci. 49:289–300.

41. Morais-Silva G, Ferreira-Santos M, Marin MT (2016): Conessine, an H3 receptor antagonist, alters behavioral and neurochemical effects of ethanol in mice. Behavioural brain research. 305:100–107.

42. Chen YW, Morganstern I, Barson JR, Hoebel BG, Leibowitz SF (2014): Differential role of D1 and D2 receptors in the perifornical lateral hypothalamus in controlling ethanol drinking and food intake: possible interaction with local orexin neurons. Alcoholism, clinical and experimental research. 38:777–786.

43. Barson JR, Leibowitz SF (2016): Hypothalamic neuropeptide signaling in alcohol addiction. Progress in neuro-psychopharmacology & biological psychiatry. 65:321–329.

44. Pelloux Y, Baunez C (2017): Targeting the subthalamic nucleus in a preclinical model of alcohol use disorder. Psychopharmacology. 234:2127–2137.

45. Zhao Y, Castellanos FX (2016): Annual Research Review: Discovery science strategies in studies of the pathophysiology of child and adolescent psychiatric disorders - promises and limitations. Journal of child psychology and psychiatry, and allied disciplines. 57:421–439.

46. Bzdok D, Yeo BTT (2017): Inference in the age of big data: Future perspectives on neuroscience. NeuroImage. 155:549–564.

47. Gaynanova I, Li G (2019): Structural learning and integrative decomposition of multi-view data. Biometrics. 75:1121–1132.

48. Zhou G, Cichocki A, Zhang Y, Mandic DP (2016): Group Component Analysis for Multiblock Data: Common and Individual Feature Extraction. IEEE Trans Neural Netw Learn Syst. 27:2426–2439.

49. Klami A, Virtanen S, Leppaaho E, Kaski S (2015): Group Factor Analysis. IEEE Trans Neural Netw Learn Syst. 26:2136–2147.

