## Supplemental Tables for "Brain Anatomical Covariation Patterns Linked to Binge Drinking and Age at First Full Drink Prior to 21 Years"

**Table S1.** Loading of brain regions in the individual thickness component related to binge drinking and AFD<21 in the HCP sample.

| ROI | Loading |  | ROI | Loading |
| --- | --- | --- | --- | --- |
| R_Temporal pole | 0.207 |  | L_Precuneus | 0.117 |
| L_Temporal pole | 0.194 |  | R_Rostral anterior cingulate | 0.117 |
| L_Transverse temporal | 0.165 |  | L_Paracentral | 0.116 |
| R_Transverse temporal | 0.159 |  | R_Parsopercularis | 0.115 |
| L_Frontal pole | 0.158 |  | R_Rostral middle frontal | 0.115 |
| R_Frontal pole | 0.153 |  | L_Inferior parietal | 0.113 |
| R_Entorhinal | 0.151 |  | R_Precuneus | 0.112 |
| R_Parsorbitalis | 0.149 |  | R_Caudal middle frontal | 0.111 |
| L_Entorhinal | 0.148 |  | R_Fusiform | 0.111 |
| L_Superior temporal | 0.145 |  | L_Isthmus cingulate | 0.111 |
| L_Middle temporal | 0.143 |  | R_Caudal anterior cingulate | 0.111 |
| R_Superior temporal | 0.140 |  | L_Insula | 0.110 |
| L_Parsorbitalis | 0.139 |  | R_Paracentral | 0.109 |
| R_Middle temporal | 0.139 |  | R_Inferior parietal | 0.107 |
| L_Superior frontal | 0.134 |  | L_Lateral occipital | 0.104 |
| L_Lateral orbitofrontal | 0.132 |  | R_Precentral | 0.103 |
| L_Parstriangularis | 0.131 |  | R_Parahippocampal | 0.101 |
| R_Bankssts | 0.130 |  | L_Caudal anterior cingulate | 0.099 |
| R_Superior frontal | 0.129 |  | L_Medial orbitofrontal | 0.098 |
| L_Fusiform | 0.128 |  | R_Lateral occipital | 0.098 |
| L_Inferior temporal | 0.127 |  | L_Posterior cingulate | 0.098 |
| R_Lateral orbitofrontal | 0.127 |  | L_Superior parietal | 0.095 |
| L_Rostral middlefrontal | 0.126 |  | R_Insula | 0.095 |
| L_Parahippocampal | 0.125 |  | L_Postcentral | 0.094 |
| L_Rostral anterior cingulate | 0.124 |  | R_Posterior cingulate | 0.092 |
| L_Supramarginal | 0.124 |  | R_Postcentral | 0.091 |
| R_Inferior temporal | 0.123 |  | R_Superior parietal | 0.089 |
| R_Medial orbitofrontal | 0.122 |  | L_Cuneus | 0.088 |
| L_Parsopercularis | 0.121 |  | R_Cuneus | 0.078 |
| R_Parstriangularis | 0.120 |  | R_Lingual | 0.078 |
| L_Caudal middle frontal | 0.120 |  | L_Pericalcarine | 0.078 |
| L_Precentral | 0.119 |  | L_Lingual | 0.077 |
| R_Supramarginal | 0.118 |  | R_Isthmus cingulate | 0.070 |
| L_Bankssts | 0.118 |  | R_Pericalcarine | 0.065 |

Abbreviations: L, left; R, right.

**Table S2.** Loading of all 221 brain features in the joint component related to AFD<21 in the HCP sample.

| Morphometry |  | ROI |  | Loading |  | Estimate | SE | P-value | P-adj |
| --- | --- | --- | --- | --- | --- | --- | --- | --- | --- |
|  |  | Joint component |  |  |  | 0.059 | 0.018 | 0.001 | 0.016 |
| SC Vol |  | BrainStem |  | 0.434 |  | -81.408 | 234.727 | 0.729 | 1 |
| SC Vol |  | L_Thalamus proper |  | 0.193 |  | 78.816 | 90.510 | 0.385 | 1 |
| GM Vol |  | L_Superior frontal |  | 0.178 |  | 499.135 | 259.035 | 0.055 | 1 |
| GM Vol |  | R_Superior frontal |  | 0.171 |  | 396.996 | 257.663 | 0.125 | 1 |
| Area |  | L_Superior frontal |  | 0.169 |  | 47.996 | 81.380 | 0.556 | 1 |
| SC Vol |  | R_Thalamus proper |  | 0.161 |  | 109.510 | 76.377 | 0.153 | 1 |
| Area |  | R_Superior frontal |  | 0.161 |  | 38.849 | 80.913 | 0.632 | 1 |
| Area |  | R_Rostral middle frontal |  | 0.144 |  | 58.630 | 79.706 | 0.463 | 1 |
| GM Vol |  | R_Rostral middle frontal |  | 0.142 |  | 449.901 | 238.236 | 0.060 | 1 |
| Area |  | L_Rostral middle frontal |  | 0.137 |  | 5.909 | 77.229 | 0.939 | 1 |
| GM Vol |  | L_Rostral middle frontal |  | 0.137 |  | 172.542 | 235.315 | 0.464 | 1 |
| Area |  | R_Inferior parietal |  | 0.125 |  | 12.878 | 80.500 | 0.873 | 1 |
| GM Vol |  | R_Inferior parietal |  | 0.124 |  | 252.435 | 236.129 | 0.286 | 1 |
| Area |  | L_Inferior parietal |  | 0.113 |  | -61.336 | 68.535 | 0.372 | 1 |
| Area |  | L_Superior parietal |  | 0.112 |  | 75.939 | 68.057 | 0.265 | 1 |
| GM Vol |  | R_Middle temporal |  | 0.111 |  | 3.979 | 156.807 | 0.980 | 1 |
| Area |  | R_Superior parietal |  | 0.109 |  | 111.454 | 70.260 | 0.114 | 1 |
| GM Vol |  | L_Inferior parietal |  | 0.109 |  | -13.005 | 202.649 | 0.949 | 1 |
| SC Vol |  | R_Putamen |  | 0.106 |  | 157.070 | 57.686 | 0.007 | 1 |
| GM Vol |  | L_Inferior temporal |  | 0.105 |  | 295.255 | 186.846 | 0.115 | 1 |
| GM Vol |  | R_Precentral |  | 0.103 |  | 339.723 | 166.586 | 0.042 | 1 |
| Area |  | L_Supramarginal |  | 0.101 |  | 27.579 | 56.565 | 0.626 | 1 |
| Area |  | R_Lateral occipital |  | 0.101 |  | 10.649 | 64.342 | 0.869 | 1 |
| Area |  | R_Precentral |  | 0.101 |  | 60.207 | 55.144 | 0.276 | 1 |
| GM Vol |  | R_Inferior temporal |  | 0.099 |  | 334.976 | 172.822 | 0.054 | 1 |
| GM Vol |  | L_Supramarginal |  | 0.099 |  | 283.881 | 171.715 | 0.099 | 1 |
| SC Vol |  | L_Putamen |  | 0.098 |  | 218.690 | 79.681 | 0.006 | 1 |
| GM Vol |  | L_Middle temporal |  | 0.098 |  | 45.376 | 151.216 | 0.764 | 1 |
| GM Vol |  | L_Superior temporal |  | 0.098 |  | 475.894 | 160.804 | 0.003 | 0.678 |
| GM Vol |  | R_Lateral occipital |  | 0.098 |  | 189.121 | 178.274 | 0.290 | 1 |
| Area |  | R_Precuneus |  | 0.096 |  | 20.679 | 53.282 | 0.698 | 1 |
| GM Vol |  | L_Superior parietal |  | 0.096 |  | 399.375 | 195.711 | 0.042 | 1 |
| GM Vol |  | R_Superior parietal |  | 0.096 |  | 427.459 | 189.905 | 0.025 | 1 |
| SC Vol |  | R_VentDC |  | 0.095 |  | -8.079 | 45.953 | 0.861 | 1 |
| SC Vol |  | L_VentDC |  | 0.094 |  | -9.317 | 45.252 | 0.837 | 1 |
| SC Vol |  | L_Hippo |  | 0.092 |  | 166.682 | 47.928 | 0.001 | 0.126 |
| GM Vol |  | R_Superior temporal |  | 0.092 |  | 322.495 | 162.427 | 0.048 | 1 |
| Area |  | L_Postcentral |  | 0.09 |  | 85.809 | 47.997 | 0.075 | 1 |
| Area |  | L_Precuneus |  | 0.09 |  | 21.325 | 46.656 | 0.648 | 1 |
| GM Vol |  | L_Precentral |  | 0.09 |  | 240.909 | 158.625 | 0.130 | 1 |
| Area |  | L_Lateral occipital |  | 0.089 |  | 43.650 | 59.942 | 0.467 | 1 |
| GM Vol |  | R_Precuneus |  | 0.089 |  | 195.015 | 138.959 | 0.162 | 1 |
| SC Vol |  | R_Hippo |  | 0.088 |  | 195.015 | 44.167 | 0.000 | 0.003 |
| SC Vol |  | R_Caudate |  | 0.087 |  | 62.652 | 51.573 | 0.225 | 1 |
| Area |  | L_Inferior temporal |  | 0.087 |  | 12.672 | 51.212 | 0.805 | 1 |
| Area |  | L_Precentral |  | 0.087 |  | 6.399 | 56.065 | 0.909 | 1 |
| GM Vol |  | R_Fusiform |  | 0.087 |  | 76.653 | 151.942 | 0.614 | 1 |
| GM Vol |  | L_Fusiform |  | 0.086 |  | -41.264 | 156.146 | 0.792 | 1 |
| GM Vol |  | L_Lateral occipital |  | 0.085 |  | 279.685 | 165.535 | 0.092 | 1 |
| GM Vol |  | R_Supramarginal |  | 0.085 |  | 2.400 | 168.613 | 0.989 | 1 |
| SC Vol |  | L_Caudate |  | 0.084 |  | 51.931 | 51.084 | 0.310 | 1 |
| Area |  | R_Supramarginal |  | 0.084 |  | -48.103 | 54.569 | 0.379 | 1 |
| Area |  | R_Inferior temporal |  | 0.083 |  | 9.329 | 47.510 | 0.844 | 1 |
| Area |  | R_Middle temporal |  | 0.083 |  | -58.430 | 40.996 | 0.155 | 1 |
| Area |  | L_Superior temporal |  | 0.082 |  | 53.378 | 43.626 | 0.222 | 1 |
| GM Vol |  | L_Postcentral |  | 0.082 |  | 362.104 | 133.833 | 0.007 | 1 |
| GM Vol |  | L_Precuneus |  | 0.082 |  | 117.548 | 133.209 | 0.378 | 1 |
| Area |  | R_Postcentral |  | 0.08 |  | 73.387 | 47.359 | 0.122 | 1 |
| Area |  | R_Fusiform |  | 0.079 |  | 2.571 | 43.073 | 0.952 | 1 |
| Area |  | L_Middle temporal |  | 0.078 |  | -33.820 | 40.242 | 0.401 | 1 |
| Area |  | L_Fusiform |  | 0.075 |  | -47.716 | 44.755 | 0.287 | 1 |
| Area |  | R_Superior temporal |  | 0.075 |  | -4.990 | 40.525 | 0.902 | 1 |
| GM Vol |  | R_Postcentral |  | 0.072 |  | 326.285 | 142.376 | 0.023 | 1 |
| Area |  | L_Lingual |  | 0.066 |  | 53.193 | 47.713 | 0.266 | 1 |
| Area |  | R_Lingual |  | 0.065 |  | 74.195 | 48.470 | 0.127 | 1 |
| GM Vol |  | R_Lateral orbitofrontal |  | 0.058 |  | 58.184 | 94.308 | 0.538 | 1 |
| GM Vol |  | L_Caudal middle frontal |  | 0.057 |  | 294.499 | 147.954 | 0.048 | 1 |
| GM Vol |  | L_Lateral orbitofrontal |  | 0.057 |  | 117.347 | 90.020 | 0.193 | 1 |
| GM Vol |  | L_Lingual |  | 0.057 |  | 293.294 | 132.281 | 0.027 | 1 |
| Area |  | L_Caudal middle frontal |  | 0.056 |  | 59.579 | 46.527 | 0.201 | 1 |
| GM Vol |  | R_Lingual |  | 0.055 |  | 310.022 | 122.228 | 0.012 | 1 |
| Area |  | R_Lateral orbitofrontal |  | 0.054 |  | -17.578 | 31.104 | 0.572 | 1 |
| GM Vol |  | R_Insula |  | 0.054 |  | 184.801 | 80.549 | 0.023 | 1 |
| Area |  | L_Lateral orbitofrontal |  | 0.052 |  | -2.592 | 28.114 | 0.927 | 1 |
| GM Vol |  | R_Caudal middle frontal |  | 0.049 |  | 102.456 | 140.187 | 0.465 | 1 |
| GM Vol |  | L_Insula |  | 0.048 |  | 169.816 | 75.298 | 0.025 | 1 |
| Area |  | R_Caudal middle frontal |  | 0.047 |  | 3.939 | 45.003 | 0.930 | 1 |
| Area |  | R_Insula |  | 0.047 |  | 18.651 | 26.125 | 0.476 | 1 |
| Area |  | L_Medial orbitofrontal |  | 0.046 |  | 63.702 | 31.809 | 0.046 | 1 |
| SC Vol |  | R_Amygdala |  | 0.043 |  | 53.861 | 20.578 | 0.009 | 1 |
| Area |  | L_Insula |  | 0.042 |  | 9.098 | 26.379 | 0.730 | 1 |
| GM Vol |  | L_Medial orbitofrontal |  | 0.042 |  | 256.406 | 83.257 | 0.002 | 0.472 |
| SC Vol |  | L_Amygdala |  | 0.04 |  | 63.138 | 18.881 | 0.001 | 0.197 |
| Area |  | R_Medial orbitofrontal |  | 0.039 |  | 6.158 | 22.896 | 0.788 | 1 |
| GM Vol |  | R_Medial orbitofrontal |  | 0.039 |  | 75.626 | 76.972 | 0.327 | 1 |
| GM Vol |  | L_Pars opercularis |  | 0.034 |  | 129.893 | 109.807 | 0.238 | 1 |
| Area |  | L_Pars opercularis |  | 0.032 |  | 2.099 | 32.700 | 0.949 | 1 |
| Area |  | R_Paracentral |  | 0.032 |  | 10.122 | 26.493 | 0.703 | 1 |
| GM Vol |  | R_Parso percularis |  | 0.032 |  | 120.245 | 96.341 | 0.213 | 1 |
| GM Vol |  | R_Paracentral |  | 0.031 |  | 79.069 | 81.893 | 0.335 | 1 |
| Area |  | R_Cuneus |  | 0.03 |  | 8.095 | 25.192 | 0.748 | 1 |
| Area |  | R_Pars opercularis |  | 0.03 |  | 20.368 | 29.571 | 0.492 | 1 |
| Area |  | R_Pericalcarine |  | 0.03 |  | -0.033 | 30.034 | 0.999 | 1 |
| Area |  | R_Posterior cingulate |  | 0.03 |  | 24.823 | 21.982 | 0.260 | 1 |
| GM Vol |  | L_Rostral anterior cingulate |  | 0.03 |  | 13.616 | 67.189 | 0.840 | 1 |
| SC Vol |  | R_Pallidum |  | 0.029 |  | 10.498 | 20.217 | 0.604 | 1 |
| Area |  | L_Pericalcarine |  | 0.029 |  | 1.281 | 27.237 | 0.963 | 1 |
| Area |  | L_Cuneus |  | 0.027 |  | 18.303 | 23.610 | 0.439 | 1 |
| Area |  | L_Posterior cingulate |  | 0.027 |  | 6.639 | 23.313 | 0.776 | 1 |
| Area |  | L_Rostral anterior cingulate |  | 0.026 |  | -5.940 | 18.083 | 0.743 | 1 |
| GM Vol |  | R_Posterior cingulate |  | 0.026 |  | 121.475 | 62.918 | 0.055 | 1 |
| Thickness |  | L_Superior temporal |  | 0.025 |  | 0.052 | 0.017 | 0.003 | 0.532 |
| Area |  | L_Paracentral |  | 0.025 |  | -1.327 | 21.399 | 0.951 | 1 |
| GM Vol |  | L_Posterior cingulate |  | 0.025 |  | 119.987 | 59.500 | 0.045 | 1 |
| GM Vol |  | R_Cuneus |  | 0.025 |  | 52.254 | 62.452 | 0.403 | 1 |
| Thickness |  | R_Superior temporal |  | 0.024 |  | 0.063 | 0.016 | 0.000 | 0.038 |
| Area |  | L_Bankssts |  | 0.024 |  | -20.109 | 19.315 | 0.299 | 1 |
| Area |  | R_Caudal anterior cingulate |  | 0.024 |  | 23.952 | 24.307 | 0.325 | 1 |
| Area |  | R_Isthmus cingulate |  | 0.024 |  | 9.840 | 18.577 | 0.597 | 1 |
| GM Vol |  | L_Bankssts |  | 0.024 |  | -13.683 | 59.728 | 0.819 | 1 |
| GM Vol |  | R_Pars triangularis |  | 0.024 |  | 194.919 | 96.161 | 0.044 | 1 |
| Thickness |  | R_Caudal anterior cingulate |  | 0.023 |  | 0.041 | 0.029 | 0.156 | 1 |
| Area |  | L_Isthmus cingulate |  | 0.023 |  | -83.308 | 52.206 | 0.112 | 1 |
| Area |  | L_Pars triangularis |  | 0.023 |  | 8.290 | 23.847 | 0.728 | 1 |
| Area |  | R_Pars triangularis |  | 0.023 |  | 30.453 | 29.742 | 0.307 | 1 |
| GM Vol |  | L_Paracentral |  | 0.023 |  | 53.550 | 68.970 | 0.438 | 1 |
| GM Vol |  | L_Pars triangularis |  | 0.023 |  | 66.934 | 78.125 | 0.392 | 1 |
| Area |  | R_Bankssts |  | 0.021 |  | 10.993 | 15.400 | 0.476 | 1 |
| GM Vol |  | L_Cuneus |  | 0.021 |  | 78.611 | 57.662 | 0.174 | 1 |
| GM Vol |  | R_Bankssts |  | 0.021 |  | 77.182 | 49.085 | 0.117 | 1 |
| GM Vol |  | R_Pericalcarine |  | 0.021 |  | 87.025 | 62.504 | 0.165 | 1 |
| GM Vol |  | R_Rostral anterior cingulate |  | 0.021 |  | 113.428 | 61.464 | 0.066 | 1 |
| SC Vol |  | L_Pallidum |  | 0.02 |  | 18.608 | 25.510 | 0.466 | 1 |
| Thickness |  | R_Transverse temporal |  | 0.02 |  | 0.088 | 0.023 | 0.000 | 0.034 |
| GM Vol |  | R_Caudal anterior cingulate |  | 0.02 |  | 111.395 | 71.696 | 0.121 | 1 |
| Area |  | R_Rostral anterior cingulate |  | 0.019 |  | 25.763 | 16.073 | 0.110 | 1 |
| GM Vol |  | L_Caudal anterior cingulate |  | 0.019 |  | 79.028 | 67.783 | 0.245 | 1 |
| GM Vol |  | L_Isthmus cingulate |  | 0.019 |  | 43.306 | 64.521 | 0.503 | 1 |
| GM Vol |  | L_Pericalcarine |  | 0.019 |  | 60.305 | 54.711 | 0.271 | 1 |
| GM Vol |  | R_Isthmus cingulate |  | 0.019 |  | 43.594 | 49.200 | 0.376 | 1 |
| Thickness |  | R_Middle temporal |  | 0.018 |  | 0.057 | 0.016 | 0.001 | 0.117 |
| Thickness |  | R_Insula |  | 0.018 |  | 0.058 | 0.017 | 0.001 | 0.168 |
| Area |  | L_Caudal anterior cingulate |  | 0.018 |  | 11.420 | 18.726 | 0.542 | 1 |
| SC Vol |  | R_Accumbens area |  | 0.017 |  | 29.655 | 11.955 | 0.014 | 1 |
| Thickness |  | R_Bankssts |  | 0.017 |  | 0.049 | 0.019 | 0.011 | 1 |
| Thickness |  | L_Inferior temporal |  | 0.016 |  | 0.055 | 0.016 | 0.001 | 0.165 |
| Thickness |  | L_Insula |  | 0.016 |  | 0.054 | 0.019 | 0.004 | 0.877 |
| Thickness |  | R_Lateral occipital |  | 0.016 |  | 0.036 | 0.013 | 0.006 | 1 |
| Area |  | R_Parahippocampal |  | 0.016 |  | 4.031 | 10.950 | 0.713 | 1 |
| GM Vol |  | R_Pars orbitalis |  | 0.016 |  | 44.378 | 45.127 | 0.326 | 1 |
| SC Vol |  | L_Accumbens area |  | 0.015 |  | 19.159 | 10.990 | 0.082 | 1 |
| Thickness |  | L_Parahippocampal |  | 0.015 |  | 0.075 | 0.034 | 0.028 | 1 |
| Thickness |  | L_Bankssts |  | 0.014 |  | 0.037 | 0.018 | 0.039 | 1 |
| Thickness |  | L_Lateral orbitofrontal |  | 0.014 |  | 0.046 | 0.016 | 0.005 | 0.979 |
| Thickness |  | L_Lingual |  | 0.014 |  | 0.039 | 0.014 | 0.008 | 1 |
| Thickness |  | L_Middle temporal |  | 0.014 |  | 0.046 | 0.018 | 0.010 | 1 |
| Thickness |  | L_Pars opercularis |  | 0.014 |  | 0.042 | 0.016 | 0.009 | 1 |
| Thickness |  | L_Postcentral |  | 0.014 |  | 0.030 | 0.012 | 0.017 | 1 |
| Thickness |  | R_Entorhinal |  | 0.014 |  | 0.088 | 0.032 | 0.007 | 1 |
| Thickness |  | R_Pars opercularis |  | 0.014 |  | 0.043 | 0.016 | 0.008 | 1 |
| Area |  | L_Parahippocampal |  | 0.014 |  | 17.793 | 12.729 | 0.163 | 1 |
| Area |  | R_Pars orbitalis |  | 0.014 |  | -7.140 | 11.235 | 0.526 | 1 |
| GM Vol |  | L_Pars orbitalis |  | 0.014 |  | 49.187 | 37.538 | 0.191 | 1 |
| Thickness |  | R_Postcentral |  | 0.013 |  | 0.028 | 0.012 | 0.023 | 1 |
| Area |  | L_Pars orbitalis |  | 0.013 |  | 3.605 | 9.439 | 0.703 | 1 |
| GM Vol |  | R_Parahippocampal |  | 0.013 |  | 59.510 | 34.733 | 0.088 | 1 |
| Thickness |  | L_Lateral occipital |  | 0.012 |  | 0.030 | 0.014 | 0.032 | 1 |
| Thickness |  | R_Inferior temporal |  | 0.012 |  | 0.074 | 0.016 | 0.000 | 0.001 |
| GM Vol |  | L_Entorhinal |  | 0.012 |  | 69.674 | 40.223 | 0.084 | 1 |
| Thickness |  | L_Fusiform |  | 0.011 |  | 0.048 | 0.015 | 0.002 | 0.396 |
| Thickness |  | R_Lingual |  | 0.011 |  | 0.032 | 0.015 | 0.039 | 1 |
| Thickness |  | R_Pars triangularis |  | 0.011 |  | 0.052 | 0.015 | 0.001 | 0.136 |
| Thickness |  | R_Temporal pole |  | 0.011 |  | 0.122 | 0.040 | 0.003 | 0.561 |
| GM Vol |  | L_Parahippocampal |  | 0.011 |  | 116.379 | 41.883 | 0.006 | 1 |
| GM Vol |  | R_Entorhinal |  | 0.011 |  | 44.724 | 43.348 | 0.303 | 1 |
| Thickness |  | L_Pars triangularis |  | 0.01 |  | 0.022 | 0.017 | 0.176 | 1 |
| Area |  | L_Entorhinal |  | 0.01 |  | 11.629 | 8.706 | 0.183 | 1 |
| Thickness |  | L_Rostral middle frontal |  | 0.009 |  | 0.031 | 0.016 | 0.046 | 1 |
| Thickness |  | R_Rostral anterior cingulate |  | 0.009 |  | 0.046 | 0.027 | 0.091 | 1 |
| Area |  | L_Transverse temporal |  | 0.009 |  | 11.114 | 8.543 | 0.194 | 1 |
| GM Vol |  | L_Temporal pole |  | 0.009 |  | 72.602 | 44.244 | 0.102 | 1 |
| GM Vol |  | L_Transverse temporal |  | 0.009 |  | 44.084 | 29.096 | 0.131 | 1 |
| Thickness |  | L_Pericalcarine |  | 0.008 |  | 0.038 | 0.014 | 0.008 | 1 |
| Thickness |  | L_Precentral |  | 0.008 |  | 0.040 | 0.015 | 0.009 | 1 |
| Thickness |  | R_Caudal middle frontal |  | 0.008 |  | 0.043 | 0.015 | 0.005 | 1 |
| Thickness |  | R_Lateral orbitofrontal |  | 0.008 |  | 0.043 | 0.016 | 0.008 | 1 |
| Thickness |  | R_Pericalcarine |  | 0.008 |  | 0.045 | 0.015 | 0.003 | 0.676 |
| Thickness |  | R_Precentral |  | 0.008 |  | 0.033 | 0.014 | 0.019 | 1 |
| Thickness |  | R_Supramarginal |  | 0.008 |  | 0.037 | 0.014 | 0.012 | 1 |
| Thickness |  | R_Frontal pole |  | 0.008 |  | 0.046 | 0.027 | 0.088 | 1 |
| Thickness |  | L_Caudal middle frontal |  | 0.007 |  | 0.033 | 0.016 | 0.043 | 1 |
| Thickness |  | L_Precuneus |  | 0.007 |  | 0.033 | 0.015 | 0.024 | 1 |
| Thickness |  | L_Supramarginal |  | 0.007 |  | 0.050 | 0.015 | 0.001 | 0.175 |
| Thickness |  | R_Fusiform |  | 0.007 |  | 0.033 | 0.014 | 0.023 | 1 |
| Thickness |  | R_Inferior parietal |  | 0.007 |  | 0.042 | 0.014 | 0.002 | 0.420 |
| Thickness |  | R_Precuneus |  | 0.007 |  | 0.031 | 0.015 | 0.037 | 1 |
| Area |  | L_Temporal pole |  | 0.007 |  | 8.508 | 7.006 | 0.226 | 1 |
| Area |  | R_Entorhinal |  | 0.007 |  | 2.601 | 9.964 | 0.794 | 1 |
| GM Vol |  | R_Transverse temporal |  | 0.007 |  | 42.783 | 25.551 | 0.095 | 1 |
| Thickness |  | L_Cuneus |  | 0.006 |  | 0.018 | 0.014 | 0.208 | 1 |
| Thickness |  | R_Cuneus |  | 0.006 |  | 0.027 | 0.014 | 0.064 | 1 |
| Thickness |  | R_Isthmus cingulate |  | 0.006 |  | 0.024 | 0.022 | 0.269 | 1 |
| Thickness |  | R_Parahippocampal |  | 0.006 |  | 0.058 | 0.027 | 0.035 | 1 |
| Thickness |  | R_Posterior cingulate |  | 0.006 |  | 0.029 | 0.019 | 0.129 | 1 |
| Area |  | R_Transverse temporal |  | 0.006 |  | 1.514 | 6.707 | 0.822 | 1 |
| GM Vol |  | R_Frontal pole |  | 0.006 |  | 42.643 | 21.914 | 0.053 | 1 |
| Thickness |  | L_Inferior parietal |  | 0.005 |  | 0.037 | 0.015 | 0.012 | 1 |
| Thickness |  | L_Medial orbitofrontal |  | 0.005 |  | 0.021 | 0.021 | 0.316 | 1 |
| Thickness |  | L_Superior parietal |  | 0.005 |  | 0.030 | 0.014 | 0.030 | 1 |
| Thickness |  | L_Temporal pole |  | 0.005 |  | 0.041 | 0.034 | 0.231 | 1 |
| Thickness |  | L_Transverse temporal |  | 0.005 |  | 0.031 | 0.024 | 0.189 | 1 |
| Thickness |  | R_Medial orbitofrontal |  | 0.005 |  | 0.031 | 0.018 | 0.093 | 1 |
| Thickness |  | R_Paracentral |  | 0.005 |  | 0.029 | 0.016 | 0.079 | 1 |
| Thickness |  | R_Rostral middle frontal |  | 0.005 |  | 0.035 | 0.015 | 0.021 | 1 |
| Area |  | R_Frontal pole |  | 0.005 |  | 6.276 | 5.302 | 0.238 | 1 |
| GM Vol |  | L_Frontal pole |  | 0.005 |  | 36.297 | 18.440 | 0.050 | 1 |
| Thickness |  | L_Isthmus cingulate |  | 0.004 |  | 0.109 | 0.042 | 0.010 | 1 |
| Thickness |  | L_Pars orbitalis |  | 0.004 |  | 0.053 | 0.019 | 0.006 | 1 |
| Thickness |  | R_Superior parietal |  | 0.004 |  | 0.027 | 0.013 | 0.033 | 1 |
| Area |  | L_Frontal pole |  | 0.004 |  | 6.694 | 4.520 | 0.140 | 1 |
| GM Vol |  | R_Temporal pole |  | 0.004 |  | 82.988 | 45.243 | 0.068 | 1 |
| Thickness |  | L_Entorhinal |  | 0.003 |  | 0.087 | 0.031 | 0.006 | 1 |
| Thickness |  | L_Rostral anterior cingulate |  | 0.003 |  | 0.027 | 0.025 | 0.284 | 1 |
| Thickness |  | L_Superior frontal |  | 0.003 |  | 0.039 | 0.018 | 0.028 | 1 |
| Thickness |  | L_Frontal pole |  | 0.003 |  | 0.043 | 0.026 | 0.107 | 1 |
| Area |  | R_Temporal pole |  | 0.003 |  | 0.669 | 7.085 | 0.925 | 1 |
| Thickness |  | R_Pars orbitalis |  | 0.002 |  | 0.045 | 0.018 | 0.014 | 1 |
| Thickness |  | L_Caudal anterior cingulate |  | 0.001 |  | 0.064 | 0.023 | 0.006 | 1 |
| Thickness |  | L_Posterior cingulate |  | 0.001 |  | 0.060 | 0.017 | 0.000 | 0.105 |
| Thickness |  | R_Superior frontal |  | 0.001 |  | 0.042 | 0.017 | 0.013 | 1 |
| Thickness |  | L_Paracentral |  | 0 |  | 0.029 | 0.017 | 0.100 | 1 |

Abbreviations: ROI, region-of-interest; L, left; R, right. SC Vol, subcortical volume; GM Vol, gray matter volume; Area, surface area; SE, standard error. P-adj., p-value after multiple comparison adjustment.

**Table S3.** Summary of binge drinking and heathy control subjects from the NKI-RS sample used in this study.

| Variables | HC | BD-AUD | BD+AUD |
| --- | --- | --- | --- |
| Number of subjects | 46 | 30 | 17 |
| Age (SD) * | 44.8 (13.4) | 31.8 (12.9) | 39.4 (13.0) |
| Sex: Female | 35 (76.1%) | 21 (70.0%) | 17 (64.7%) |
| Race: White * | 33 (71.7%) | 20 (66.7%) | 17 (100%) |
| Handedness (SD) | 76.3 (28.9) | 68.0 (45.6) | 87.6 (11.3) |
| Intelligence: WASI-II (SD) * | 105.0 (11.0) | 96.7 (12.6) | 98.3 (13.2) |
| Total intracranial volume cm^3^ (SD) | 1433.3 (162.3) | 1464.9 (184.2) | 1483.6 (161.1) |
| Age at first full drink (SD) | 17.0 (3.5) | 16.7 (2.0) | 15.2 (2.0) |

HC, healthy controls; BD-AUD, BD without AUD, BD+AUD, BD with AUD. WASI-II, Wechsler Abbreviated Scale of Intelligence-II. SD, standard deviation. * indicates a significant difference at 0.05 level (without multiple comparison adjustment).

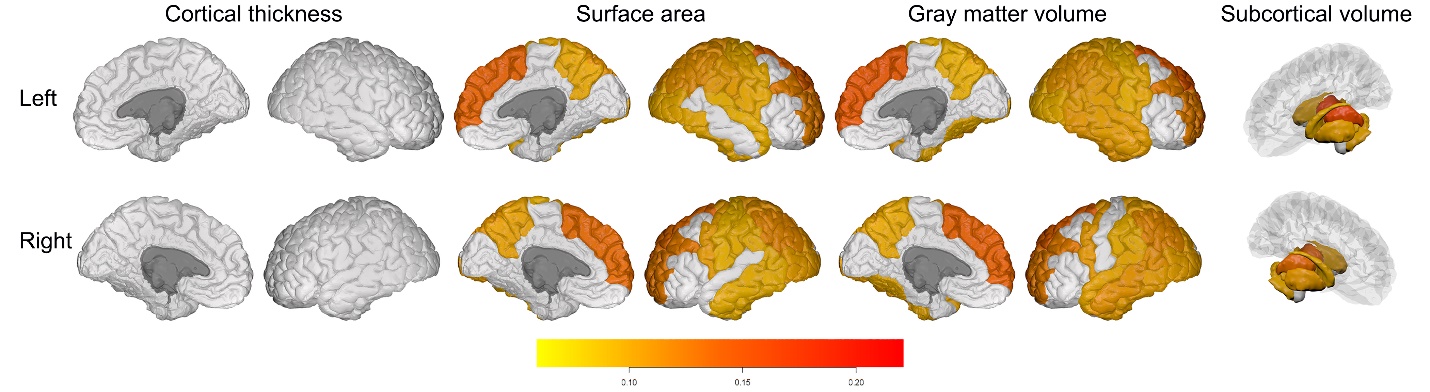

**Figure S1.** Brain regions in the joint component with loadings larger than 0.08.
